# MicroCT optimisation for imaging fascicular anatomy in peripheral nerves

**DOI:** 10.1101/818237

**Authors:** Nicole Thompson, Enrico Ravagli, Svetlana Mastitskaya, Francesco Iacoviello, Kirill Aristovich, Justin Perkins, Paul R Shearing, David Holder

## Abstract

Vagus nerve stimulation (VNS) is a promising therapy for treatment of various conditions resistant to standard therapeutics. However, due to the lack of understanding of the fascicular organisation of the vagus nerve, VNS leads to unwanted off-target effects. Micro-computed tomography (microCT) can be used to trace fascicles from periphery and image fascicular anatomy. In this work we optimised the microCT protocol of the rat sciatic and subsequent pig vagus nerves.

After differential staining, the optimal staining time was selected and scanning parameters were altered in subsequent scans. Scans were reconstructed, visualised in ImageJ and fascicles segmented with a custom algorithm in Matlab to determine ultimate parameters for tracking of the nerve. Successful segmentation for tracking of individual fascicles was achieved after 24 hours and 120 hours of staining with Lugol’s solution (1% total iodine) for rat sciatic and pig vagus nerves, respectively, and the following scanning parameters: 4 µm voxel size, 35 kVp energy, 114 µA current, 4 W power, 0.25 fps in 4 s exposure time, 3176 projections and a molybdenum target.

The optimised microCT protocol allows for segmentation and tracking of the fascicles within the nerve. This will be used to scan the full length of the pig, and possibly, the human vagus nerves. The resulting segmentation map of the functional anatomical organisation of the vagus nerve will enable selective VNS ultimately allowing for the avoidance of the off-target effects and improving its therapeutic efficacy.

## 1. Introduction

Peripheral nerves comprise hundreds or thousands of nerve fibres. These are arranged into bundles, termed fascicles, separated by perineurium and interfascicular epineurium. This is composed of epithelioid myofibroblasts which have epithelioid and myofibroblastoid properties including tight junctions, gap junctions, external laminae and contractility. The tight junctions provide selective barriers to chemical substances. Their anatomy is relatively little studied. It is fairly well documented for major nerves in the somatic nervous system, where the fascicles may be shown to have a somatotopic connection to muscles and skin (Sunderland, 1978). However, their functional anatomy in the autonomic nervous system (ANS) is almost completely unknown.

Determining this in the ANS has recently become relevant because of the development of Electroceuticals, treatment of endocrine, autoimmune and other diseases by electrical stimulation of autonomic nerves (Bonaz et al., 2018; Breit et al., 2018; Browning et al., 2017; Koopman et al., 2016). At present, this is achieved by electrical stimulation of the entirety of the nerve. As most nerves supply multiple organs and functions, this may lead to unwanted off-target and insufficient therapeutic effects. This is especially relevant to stimulation of the cervical vagus nerve, which is a convenient and surgically accessible target in the neck, known to innervate the heart, lungs and abdominal viscera (Thompson et al., 2019). The left and right cervical vagus nerves comprise a total of 10 to 15 fascicles in the adult human but their functional anatomy is unknown (Verlinden et al., 2016). This work arose out of a desire to minimise side effects by selective stimulation of relevant fascicles in the cervical vagus nerve. To identify the function of fascicles observed on histological section, we required a method to trace the fascicles from the neck to their branch insertion near supplied organs; this was intended to provide an independent validation for the new method of Electrical Impedance Tomography, which enables imaging of fascicular compound action potential activity using a cylindrical nerve cuff (Aristovich et al., 2019). MicroCT is an established tissue imaging method which has been applied for imaging of soft tissue and peripheral nerve after staining with a contrast agent. In this paper, we have developed and evaluated an optimised protocol for microCT imaging of peripheral nerves which enables segmentation and tracking of fascicles within the nerve whilst allowing for subsequent histology and validation.

### 1.1 Background

#### 1.1.1 Peripheral nervous system

The nervous system comprises the central nervous system (CNS), which includes the brain and spinal cord, and peripheral nervous system (PNS), which includes all nerves and ganglia lying outside the brain and spinal cord. The ANS is a division of the PNS. It comprises visceral afferent (sensory) and visceral efferent fibres; the latter are either parasympathetic or sympathetic (Amann and Constantinescu, 1990). The vagus nerve is one of the principal cranial nerves which contributes to the parasympathetic division; it plays a role in regulatory and homeostatic reparative systems responsible for rest-and-digest activity stimulation of the body (Felten et al., 2016; McCorry, 2007; Rea, 2014; Sibilla and Agarwal, 2018).

#### 1.1.2 MicroCT

Micro-computed tomography (microCT) provides rapid visualisation with a spatial resolution of a maximum 4 µm in three dimensions with little distortion of the sample and minimal artefacts (Stauber and Müller, 2008) which is superior to the golden standard method of histology (Chatterjee, 2014; McInnes, 2005). The process of histology results in destruction of the sample, and if 3D visualisation is desired, reconstruction of serial section images is required – a process very laborious even with computer and semi-automated assistance (Metscher, 2009).

The microCT equipment comprises components including an X-ray tube, radiation filter, sample stand and the phosphor-detector (Boerckel et al., 2014). Laboratory-based microCT instruments construct 3D volumes by computational processing of a large number of 2D radiographic projections taken at sequential angles of a rotating specimen using penetrating X-rays (O’Sullivan et al., 2018; Shearer et al., 2016).

Currently, CT and microCT are standard tools for the visualisation of bone structure. However, the imaging of biological soft tissue, limited by the low intrinsic X-ray contrast of non-mineralised tissues, has increased in the recent years with methods incorporating the use of contrast enhancement agents such as osmium (Johnson et al., 2006), reduced silver (Mizutani et al., 2007), resin perfusion (Wirkner and Prendini, 2007; Wirkner and Richter, 2004), and iodine (de Crespigny et al., 2008; Degenhardt et al., 2010; Gignac and Kley, 2014; Heimel et al., 2019; Jeffery et al., 2011; Metscher, 2009; Yan et al., 2017). It has been successfully applied to tendons (Kalson et al., 2012), ligaments (Shearer et al., 2014) and nerves (Yan et al., 2017; Zhu et al., 2016). However, there are some shortcomings of the method used hitherto. Currently, they require phase contrast scanners which may be inaccessible and time-limited and are expensive. The imaging data obtained requires considerable CPU and GPU memory in addition to the use of supercomputers (Gong et al., 2016; Yan et al., 2017). Long pre-processing procedures are needed that may allow movement of the specimen during the scan or cause destruction of the sample, so that subsequent validation with histology or other techniques, such as neural tracers, is not possible. Finally, most phase-contrast scanners allow for only a small amount of specimen to be imaged at a time at the desired resolution. This is particularly relevant for the purpose of tracing anatomical projections of the nerve and tracking individual fascicles, which require scanning of the nerve in its entirety.

To visualise soft tissue details, an X-ray contrast agent is required with different binding affinities for the tissue likely to be encountered. Iodine provides suitable soft tissue contrast in X-ray imaging (Jeffery et al., 2011; Pauwels et al., 2013) by binding to lipids and glycogen within soft tissue (Heimel et al., 2019). Soft tissues of different densities absorb iodine with different efficiency, creating a clear gradient in attenuation of X-rays. Within the nerve, it provides suitable contrast to differentiate between the fascicles, the interfascicular epineurium (connective tissue), adipocytes and the surrounding air or mount holding the nerve (Gignac et al., 2016).

#### 1.1.3 Vagus nerve stimulation and off-target effects

In order to avoid the side effects that occur with pharmacological and surgical therapies, (de Boer et al., 2013; Dunlop, 1969; Mishra, 2017; Zimmerman, 1999), the new field of bioelectronics medicine, Electroceuticals, was born. This is undertaken by electrical stimulation of autonomic nerves (Famm et al., 2013; Kollewe, 2017; Mishra, 2017). A prime target for intervention is the cervical vagus nerve (Blount, 2015; Ekmekçi and Kaptan, 2017; Guiraud et al., 2016; Koopman et al., 2016; Pečlin and Rozman, 2014; Smucny et al., 2015), as it innervates the majority of the visceral organs and muscles (Câmara and Griessenauer, 2015; Rea, 2014). However, surprisingly, the organisation of fascicles within the nerve remains almost completely unknown. Electrical stimulation of the cervical vagus nerve, a procedure known as vagus nerve stimulation (VNS), has been successfully used to reduce depression, arthritis, the frequency of epileptic seizures and improve outcomes of heart failure (Binnie, 2000; De Ferrari and Schwartz, 2011; Klein and Ferrari, 2010; Ripplinger, 2017). At the moment, current practice is that the entire nerve is activated or suppressed. This leads to off-target side effects as organs other than those intended are stimulated (Ripplinger, 2017).

### 1.2 Purpose

#### 1.2.1 Rationale

In the somatic nervous system, fascicle functional anatomy appears to follow expected principles and is somatotopically organised. The organisational principles in the ANS are unknown but it appears possible that the fascicles in complex nerves, such as the vagus, are organotopic. If this is the case, then determination of the anatomical connections of fascicles by microCT could enable production of a neuroanatomical map of the functional anatomy of the cervical vagus nerve and so allow for accurate selective VNS. This could be helpful not only in avoiding side effects but also in improving the efficacy of VNS overall by providing better understanding of the innervation of vagal fibres to both peripheral organs and originating brain regions. Firstly, however, an optimised method needed to be developed in order to image nerve fascicles efficiently and accurately over the full length of the nerve. This would allow for segmentation of the fascicles from the 3D microCT images from the branches to the central region. We also hoped to preserve the nerve samples to be available for subsequent histology for validation.

#### 1.2.2 Main purpose

The ultimate purpose for this study was to improve and optimise a simple and optimised protocol for microCT imaging of the fascicles within nerves that subsequently allows for successful segmentation and thus the development of a neuroanatomical map. The aim of the study was to answer the following questions:

a. What is the optimal protocol in terms of:
  i. Iodine staining time?
  ii. MicroCT scanning parameters?
b. Does it work:
  i. At 2D level in distinguishing fascicles?
  ii. Over the length of the nerve for 3D tracing?

### 1.3 Experimental design

In this work, a fully optimised protocol for scanning and imaging the fascicles within peripheral nerves was developed.

The overall aim was to develop a method suitable for use in tracing fascicle function in the human. In this study, we refined the method using the rat sciatic and pig vagus nerves. The rat is widely used in medical research (Iannaccone and Jacob, 2009) with its sciatic nerve being a well-established model for studies of peripheral nerves (Kaplan et al., 2015). Rat sciatic nerves were used in our experiments for technical integration, development and refinement of the imaging methods. The rat vagus nerve is monofascicular and too small, therefore not suitable for the purposes of our project. The pig vagus nerve is a preclinical model for VNS (Wolthuis et al., 2016). The small but well-studied rat sciatic nerve model allowed for development of the technique in end organ or function tracking and the pig vagus nerve, across a similar length, validated the protocol as a viable method in a more similar model to the desired human vagus nerve.

Lugol’s iodine solution, of the many different contrast agents available, was selected as the contrast agent of choice due to its ease of handling, lack of toxicity, cost-effectiveness and, most importantly, differential affinities for major types of soft tissues (Gignac et al., 2016; Metscher, 2009).

There are two main methods for sample preparation previously used in similar studies; including freeze-drying (Yan et al., 2017) and paraffin embedding (Scott et al., 2015). These were both tried and discarded. Freeze-drying is a procedure used as a pre-processing method for similar purposes; however, it adds a significant amount of time (4 days) to the already time-consuming pre-processing, and produces artefacts, such as air pockets, visible during scanning. On the other hand, during the processing step of paraffin embedding, the Lugol’s solution is washed out. This, together with suspension of the nerve in the more radiopaque paraffin wax, decreases the contrast required to image the nerve and its fascicles successfully. Both of these pre-processing methods destroy the nerve sample, thus not allowing for subsequent histology. The final utilised method was fixation, Lugol’s staining and placement of specimen within cling film for scanning. This approach maintains moisture within the nerve and reduces shrinkage during the scan. The Lugol’s iodine solution can easily be soaked out and the nerve then prepared for histology.

Prior to any Matlab processing, neighbourhood connections were performed on raw median filtered data files and segmentation of fascicles was attempted as test segmentations to show feasibility as proof of concept. This showed that quality is high enough for segmentation but the algorithm in Matlab, above, was used in conjunction with Seg3D as the final step as a superior method of segmentation.

Histology of multiple sections has been a golden standard for accurate visualisation of soft tissues and nerves, specifically for neuroanatomical tracing. However, it is time-consuming and prone to mechanical artefacts. MicroCT was chosen as a non-invasive technique that could provide 3D reconstruction of the full nerve scanned and sufficient resolution for tracing fascicles. It could be free of mechanical artefacts present in histology, and save both time and labour.

MicroCT scans were undertaken with nerves stained for different lengths of time and with differing scan parameters, including the energy. They were evaluated according to a distinguishability factor and signal-to-noise ratio (SNR) criteria allowing for determination of the protocol allowing for the most suitable segmentation and tracking. The findings were validated by comparison to histological sections with H&E. Staining was varied from 0 to 10 days in 11 rat sciatic nerves, and for 1, 3, 5, 7 and 13 days in five pig vagus nerves (one nerve for each different length of staining). MicroCT parameters were varied for the scanning of five rat sciatic nerves and testing reproducibility and use of the protocol was performed with 12 rat sciatic and three pig vagus nerves.

## 2. Methods

### 2.1 Nerve samples

Rat sciatic and pig cervical vagus nerve samples were dissected from Sprague-Dawley adult male rats, weighing 400–550g, and domestic female pigs, weighing 60-70kg, respectively, subsequent to euthanasia (ethically approved by the UK Home Office and performed in accordance with its regulations, as outlined in the Animals (Scientific Procedures) Act 1986). Nerves were then placed in neutral buffered formalin (NBF) (10%) (Sigma Aldrich HT501128) to allow for fixation.

### 2.2 Optimisation of contrast staining

#### 2.2.1 Pre-processing

##### Rat sciatic nerve

Prior to the microCT scan, 1 cm lengths of the proximal common region of the rat sciatic nerve in the thigh were cut and placed into a tube of 10 ml Lugol’s solution (total iodine 1%; 0.74% KI, 0.37% I) (Sigma Aldrich L6141) for 0 to 10 days (a total of 11 samples; one sample for each number of days). The optimal staining time was determined for 1 cm lengths of the nerve, and then applied to the full rat sciatic nerve (4 cm).

##### Pig vagus nerve

The cervical vagus nerve was sectioned into 4 cm lengths and returned to formalin (neutral buffered, 10%, Sigma Aldrich HT501128-4L) until required for staining. Prior to the microCT scan, the nerve was placed into a tube of 14 ml Lugol’s solution for 1, 3, 5, 7 and 13 days (a total of 5 samples; one sample for each number of days listed).

To achieve the optimal contrast for fascicles compared to the rest of the nerve tissue within the microCT scan, nerves were stained for the varying number of days and resulting scans compared. For staining of 4 days or more, the Lugol’s solution was refreshed halfway through the total number of days of staining to allow for resaturation with iodine.

On the day of the microCT scan, the nerve was removed from the tube and blotted dry on paper towel to remove any excess of Lugol’s solution. Nerves were then placed in a tube for vertical alignment and mounted to the sample holder in the scanner. Movement artefacts were present within the first few scans. Subsequent nerves were then first placed into a piece of cling film (8 cm × 3.5 cm) (Tesco, United Kingdom) and wrapped up to seal the nerve and thus retain moisture during the scan as to avoid shrinkage of the nerve tissue. Sutures were tied around the nerve and cling film on each end of the nerve to maintain the position of the nerve within the cling film over time during the scan and to pull the nerve into a taut position within the sponge and mount. Then, the wrapped nerve was placed within a slit cut into a sponge (7 cm × 4 cm × 1.5 cm), the sponge curled into a cylinder (diameter of ∼1 cm) and secured with tape (Transpore™, 3M, United Kingdom) to remain in shape. With a pair of forceps, the sponge cylinder was inserted into the 3D printed mount and, using the sutures, the nerve was pulled straight and aligned with centre within the sponge cylinder. The mount was sealed with the lid.

#### 2.2.2 MicroCT scanning

The microCT scanner (Nikon XT H 225, Nikon Metrology, Tring, UK) was homed and then conditioned at 200 kVp for 10 minutes before scanning. The rat sciatic nerves were scanned with the parameters in the following table (Table 1). The initial parameters used were roughly based on previous studies for soft tissues (du Plessis et al., 2017; Metscher, 2009; Senter-Zapata et al., 2016; Yan et al., 2017). The same parameters were used for all scans of the rat sciatic nerves stained for different lengths of time as to allow the differentiation in the contrast in the scans to be attributed only to the staining time. The differentially stained pig vagus nerves were scanned at the optimal parameters chosen for rat sciatic nerve following analysis (Section 2.5) of scans at differential parameters (Section 2.3).

**Table 1.**
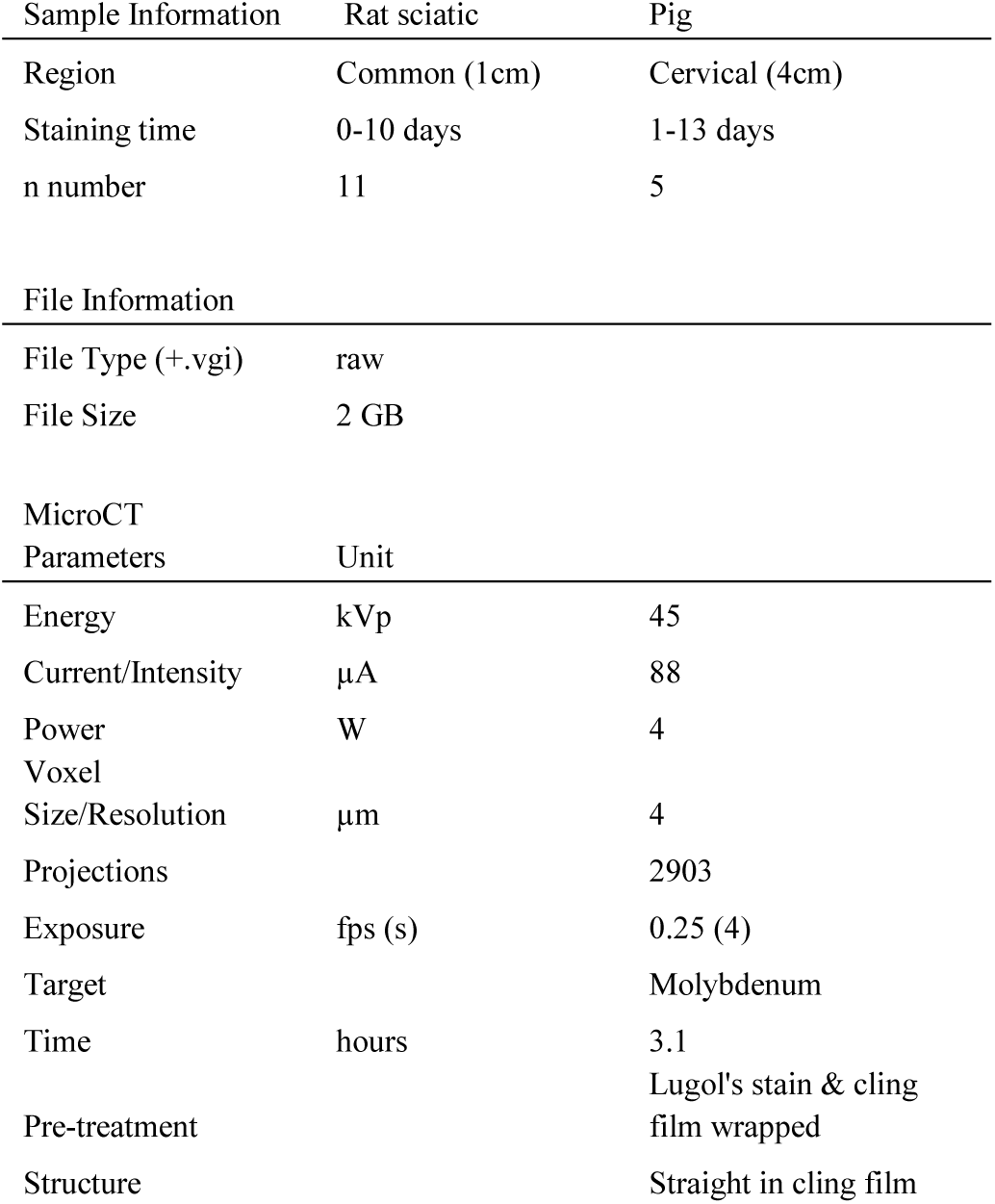
Scanning parameters to test optimal staining time of nerves.

### 2.3 Optimisation of scanning parameters

#### 2.3.1 Pre-processing

Pre-processing was performed as before (Section 2.2.1) with rat sciatic nerves being stained for 1 day and pig vagus for 5 days.

#### 2.3.2 MicroCT scanning

To optimise the scanning parameters for highest image quality and contrast, several parameters were tested for optimal value across the scans of five rat sciatic nerves (Table 2). All five scans used a molybdenum target, a power of 4 W, 3176 projections and a resolution with isotropic voxel size of 4 µm. The microCT scanner (Nikon XT H 225) was homed and then conditioned prior to scanning as described above.

**Table 2.** Differential scanning parameters for five rat sciatic nerve samples (a-e) to test optimal scanning protocol.

| Nerve Sample | Energy | Current | Exposure |
| --- | --- | --- | --- |
| | kVp | $\mu\text{A}$ | fps (s) |
| a | 45 | 88 | 0.25 (4) |
| b | 65 | 62 | 0.25 (4) |
| c | 35 | 114 | 0.25 (4) |
| d | 45 | 88 | 1 (1) |
| e | 35 | 114 | 0.5 (2) |

### 2.4 Scan reconstruction

Scans were reconstructed in CT Pro 3D (Nikon’s software for reconstructing CT data generated by Nikon Metrology, Tring, UK). Centre of rotation was calculated manually with dual slice selection. Beam hardening correction was performed with a preset of 2 and coefficient of 0.0. The reconstructions were saved as 16 bit volumes and triple TIFF 16-bit image stack files allowing for subsequent image analysis and segmentation in various software.

### 2.5 Image analysis

Reconstructed microCT scan images were analysed in ImageJ (Schindelin et al., 2012) in the XY plane to view the cross-section of the nerve. The vertical alignment of the nerve was positioned so that the cross-sectional plane is viewed in the XY stack and the longitudinal plane is viewed in the XZ and YZ stacks. This allowed for validation of the scanning protocol, direction of the nerve, and visual analysis of the quality of the image and the distinguishability of the soft tissues - specifically the identification of the fascicles known to exist within the nerve.

#### Differential staining

The greyscale intensity of the fascicle, interfascicular epineurium, and the sponge/air was calculated for each scan of nerves stained for different lengths of time. A straight line, with a length of approximately 100 pixels, was drawn starting within a fascicle, continuing through the interfascicular epineurium and ending in the sponge/air. The Plot Profile function, from the Analyze menu, was used. This function produces a two-dimensional graph of the greyscale intensities of pixels along the line within the image. The x-axis depicts the distance of the pixels along the line and the y-axis represents the grey value or intensity of the pixels. The grey values for each soft tissue type were then analysed and compared. The distinguishability value (d) was determined for each scan: the sum of the permutative differences between the absolute mean values of the three tissue types (fascicles, interfascicular epineurium and adipocytes) was calculated and maximised.

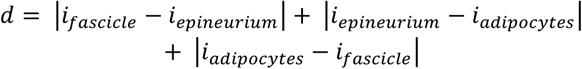

The highest value depicts the staining time that results in and provides the greatest difference in greyscale between the three tissue types. Thus, should result in the optimal segmentation of the fascicles from both the interfascicular epineurium and adipocytes allowing for tracking of the fascicles.

#### Differential parameters

The signal-to-noise ratio (SNR) of fascicle to background was calculated for each scan. A random block of sample pixels/greyscale values were obtained from both a fascicle and the background of the microCT scan images for which the average and the standard deviation of the pixel data was calculated, respectively.

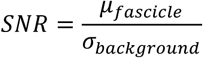

The highest SNR depicts the greatest difference between the signal and the background noise. Therefore, the highest SNR results from the microCT images that have the greatest signal from the fascicles – the object of interest for segmentation in this case.

### 2.6 Histology for validation

After dissection from the animal, the nerve was placed into neutral buffered formalin for fixation for approximately two to five days. Thereafter, nerves were embedded in paraffin, sectioned at 4 µm, and stained with Haemotoxylin and Eosin (H&E, a routine stain used to demonstrate the general morphology of tissue) (Sheehan and Hrapchak, 1987). Subsequent microscopy analysis of the tissue slides to identify histopathological features was performed. The images of the H&E sections were then compared to the corresponding slice in the microCT scan of the same nerve for comparison and validation.

The diameters of the fascicles and the full nerve in the microCT scans and the corresponding histology slice were calculated and compared to determine the percentage accuracy of the nerve and fascicle diameter measured in the microCT scan in relation to the histology image. The freehand tool in ImageJ was used to isolate the corresponding fascicles in both the microCT scan slice and the histology section, for which the area was calculated using the Measure tool and the scale of the image. The diameters were then measured for each fascicle and the full nerve, and the percentage difference between the histology and microCT images was calculated.

### 2.7 Computerised 3D reconstruction and fascicle tracking

The chosen optimal staining and scanning parameters were used for the scanning of multiple nerves (n=12 rat sciatic nerves, n=3 pig vagus nerves). The 4cm length of nerve was scanned by setting up a program of scans with an overlap region of 20%. Computerised 3D reconstruction and fascicle tracking was then performed on the reconstructed scans.

The volume data was post-processed in Matlab R2018b (Mathworks, Natick, Massachusetts, USA). The slices on the extremities affected by the common cone-beam artefacts were removed as to not interfere with further post-processing and segmentation. Histogram stretching and rescaling was performed to get rid of irrelevant pixels outside of the histogram range of the volume of interest (VOI; in this case, the fascicles). A 3D volume median filter was applied with a 3×3×3 kernel size. Convolution with a short longitudinal kernel was performed across the entire stack with a kernel of 1×1×5. The threshold for binarization was then determined to specify the region of greyscale values consisting of the fascicles; thus a volume of the fascicles was produced.

Subsequently, NIFTI files were formed to be compatible in file type and size for use in Seg3D (http://www.seg3d.org, NIH Centre for Integrative Biomedical Computing, SCI Institute, University of Utah, USA), a free volume segmentation and processing tool. The neighbourhood connected filter, that marks all the pixels within the same data range as your selected seed points, was performed and fascicles segmented. Where appropriate or required, smoothing by dilation and erosion was performed as a test for more accurate segmentation across regions of air pockets or uncharted change in greyscale, i.e. insufficient contrast between the VOI and surrounding medium despite being able to distinguish structures by eye. Completed segmentations were compared to full stack of XY slices to confirm branching regions and plexuses that may be assumed to be unsuccessful segmentation and boundary gain.

Tracing of the fascicles was deemed to be acceptable if tracing of the three main fascicles (tibial, peroneal and sural) within the rat sciatic nerve, and 10 fascicles in the pig vagus nerve, over the length of the scan was possible without loss or gain of boundary. Boundary loss is defined as the discontinuation of neighbourhood connections around more than half of the diameter of the fascicle. Boundary gain is defined as the segmentation of more than one fascicle at a time due to low resolution and loss of greyscale contrast unless fascicles are connected by a visible plexus within 3 mm of nerve. Also pertaining to boundary gain, is the unwanted segmentation of the fascicles along with adipocytes, blood vessels or interfascicular epineurium.

## 3. Results

### 3.1 Sample numbers

#### Rat sciatic nerves

differential staining (1 cm): n=11; differential parameters (1 cm): n=5; reproducibility (4 cm): n=12.

#### Pig vagus nerves

differential staining (4 cm): n=5; reproducibility (4 cm): n=3.

### 3.2 Optimisation of contrast staining

Visual inspection of the representative slices from each scan of rat sciatic nerves showed clear differentiation of the fascicles from the rest of the nerve (Figure 3), owing to saturation of the nerve with Lugol’s solution. The clearest distinction was visible for one day of staining. Inspection of the pig vagus nerve scans revealed the extent of staining varied evidently between the different days of staining; with fascicles and adipocytes not clearly visible in each scan (Figure 4). Five days of staining visibly showed the clearest distinction between soft tissue types.

**Figure 1.**
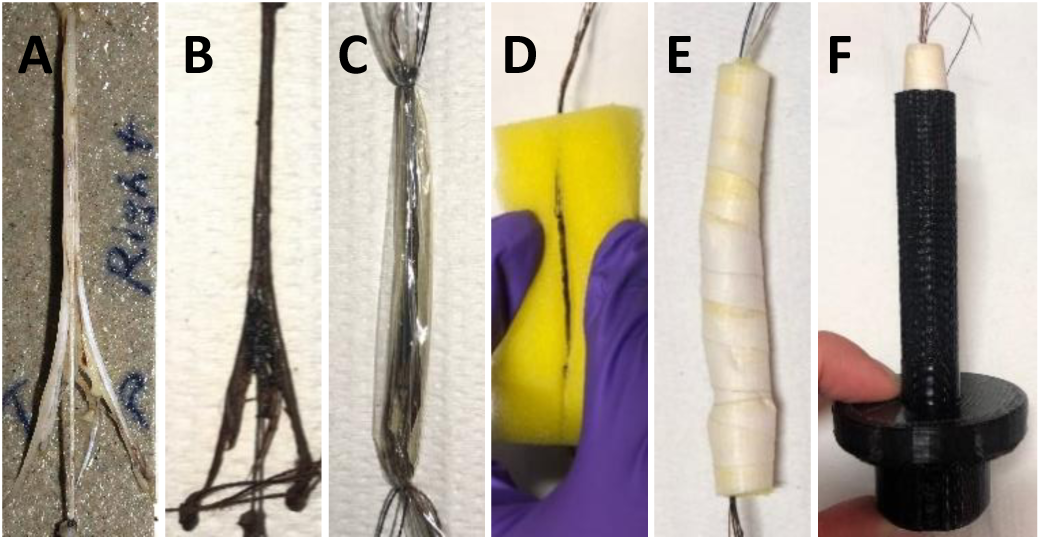
The rat sciatic nerve A. after fixation and before staining, B. after staining, C. wrapped in cling film and, D. placed in sponge. E. Sponge with nerve coiled up and, F. placed within 3D printed mount.

**Figure 2.**
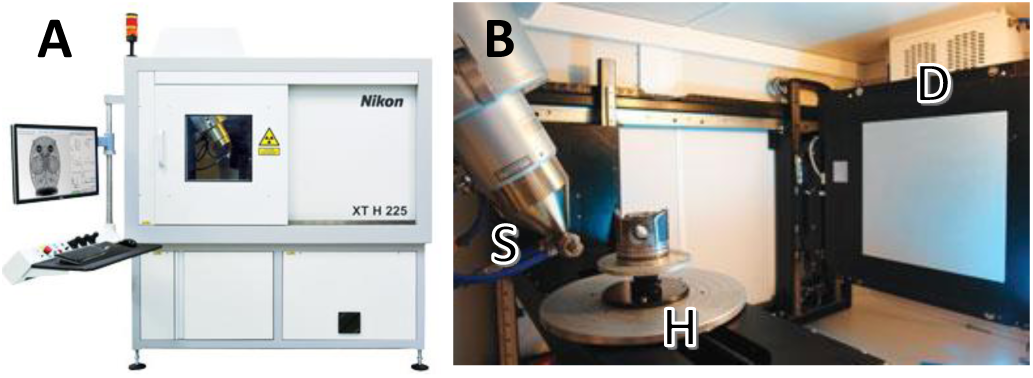
A. The Nikon XT H 225 microCT scanning machine (*“XT H 225* | *Computed Tomography* | *X-ray and CT inspection* | *Nikon Metrology,” n.d*.)., B. The internal setup of the scanner’s principal components including the open-tube X-ray source (S), rotating specimen holder (H), and the detector (D) (1620 PerkinElmer, 2K x 2K, 200 microns pitch).

**Figure 3.**
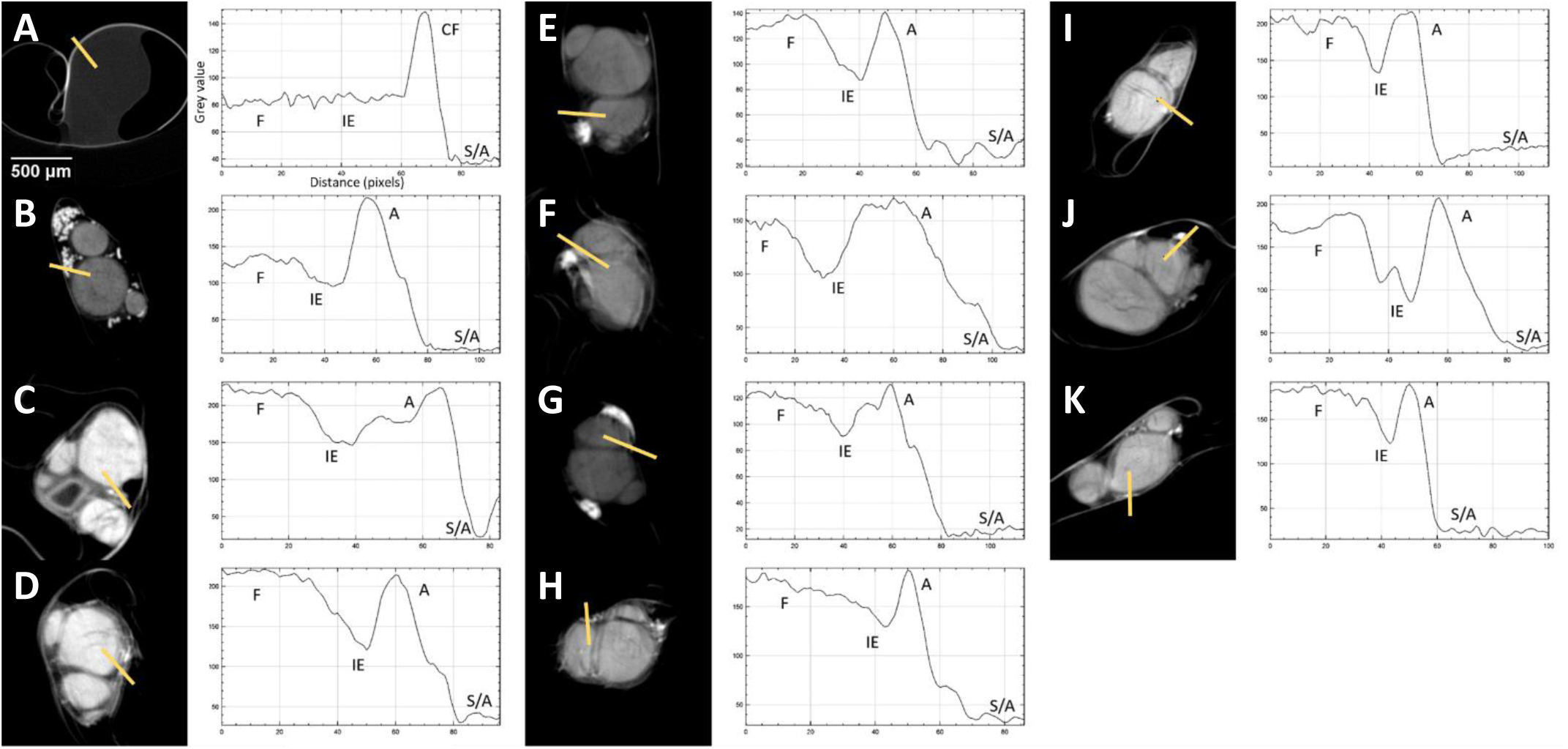
Representative XY plane slicefrom each scan with corresponding grey value graph across the distance of the line drawn through the four ROI’s with A-K being nerves scanned for 0-10 days, respectively. F = fascicle, IE = interfascicular epineurium, CF = cling film, A = adipocytes, S/A = sponge/air.

**Figure 4.**
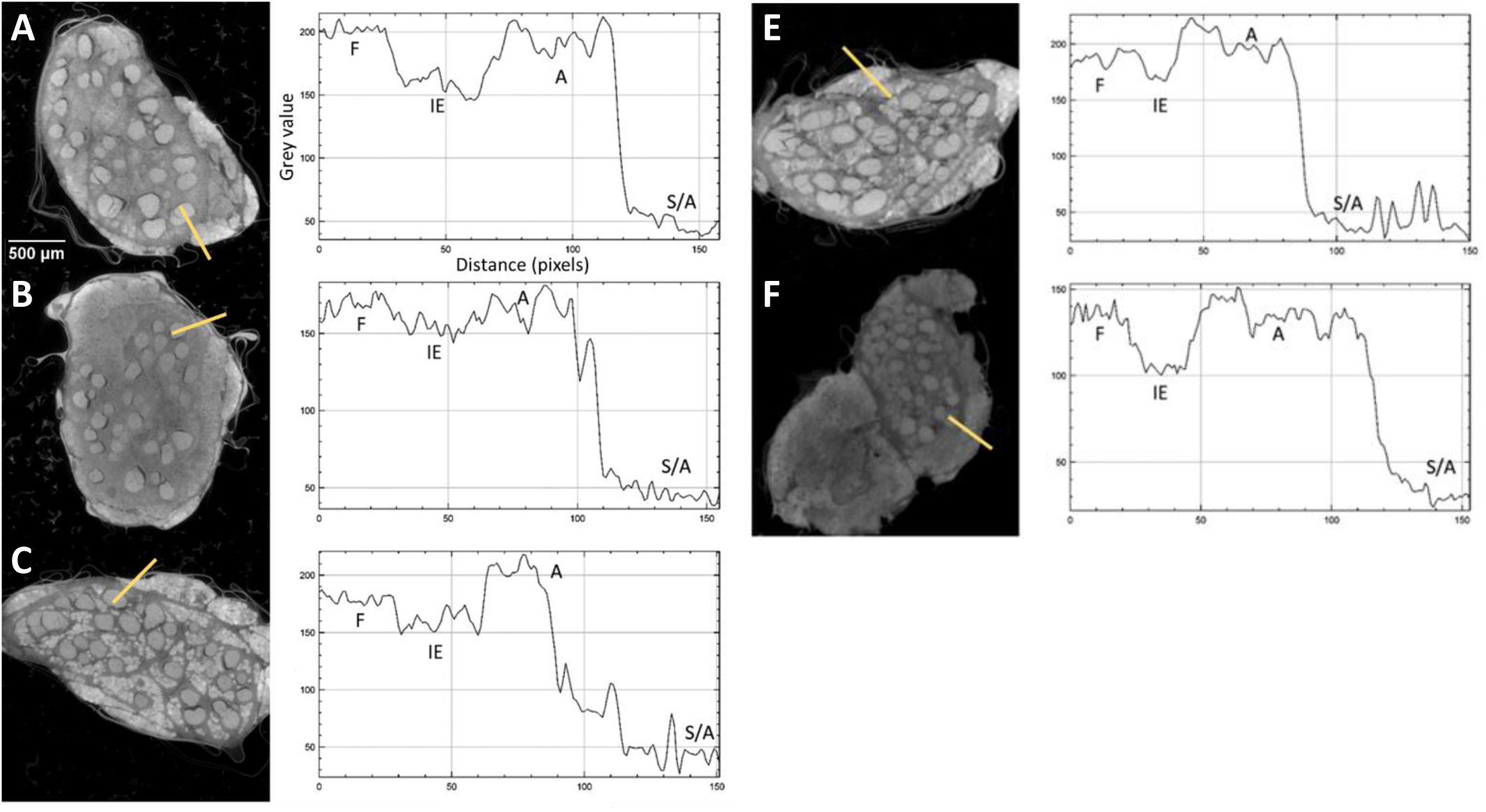
Representative XY plane slice from each scan with corresponding grey value graph across the distance of the line drawn through the four ROI’s with A-E being nerves scanned for 1, 3, 5, 7 and 13 days, respectively. F = fascicle, IE = interfascicular epineurium, CF = cling film, A = adipocytes, S/A = sponge/air.

Movement artefacts were present during some scans, however, grey values could still be calculated. With a greyscale range of 0 to 1, the average of the mean grey values of all the days of staining for the adipocytes and sponge/air in the rat sciatic nerve scans was 0.92±0.03 and 0.18±0.04, respectively (Figure 5). On the other hand, the deviation between the grey values for the fascicles and interfascicular epineurium was 0.1 and 0.09, respectively. This larger deviation is as result of one of the samples having a greater difference in mean grey values for the four different ROI. To corroborate this, the distinguishability values (d) were calculated for each scan of rat sciatic (Figure 6) and pig vagus (Figure 7) nerves (rat sciatic: n=11; pig vagus: n=5).

**Figure 5.**
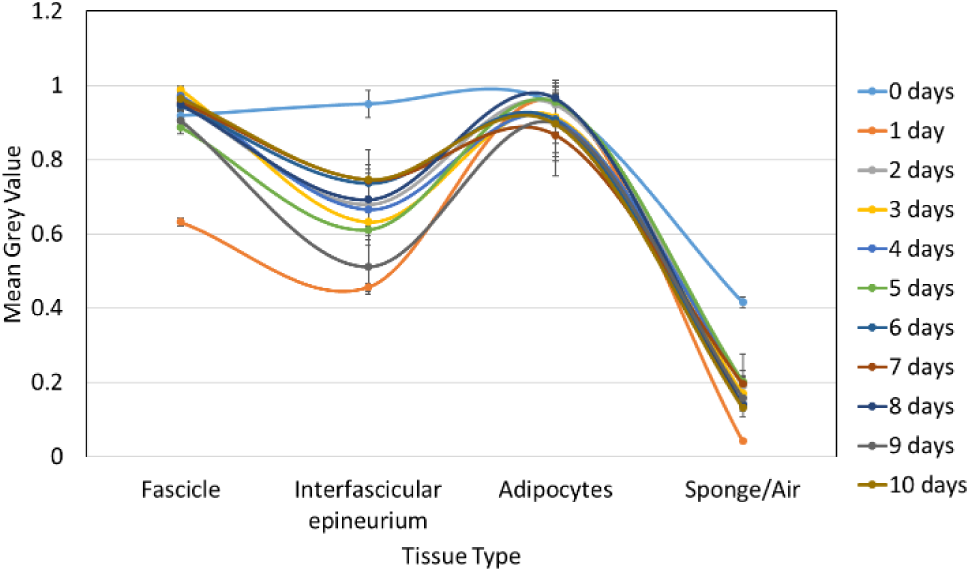
Mean grey value of the different tissue types for each rat sciatic nerve scanned after staining for a various number of days.

**Figure 6.**
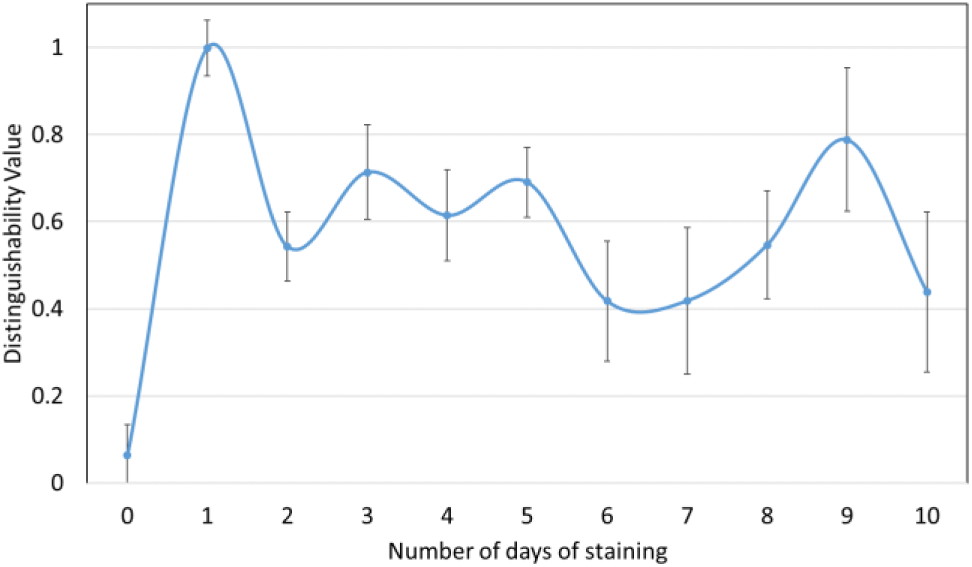
Distinguishability value for each rat sciatic nerve scanned after a different number of days of staining.

**Figure 7.**
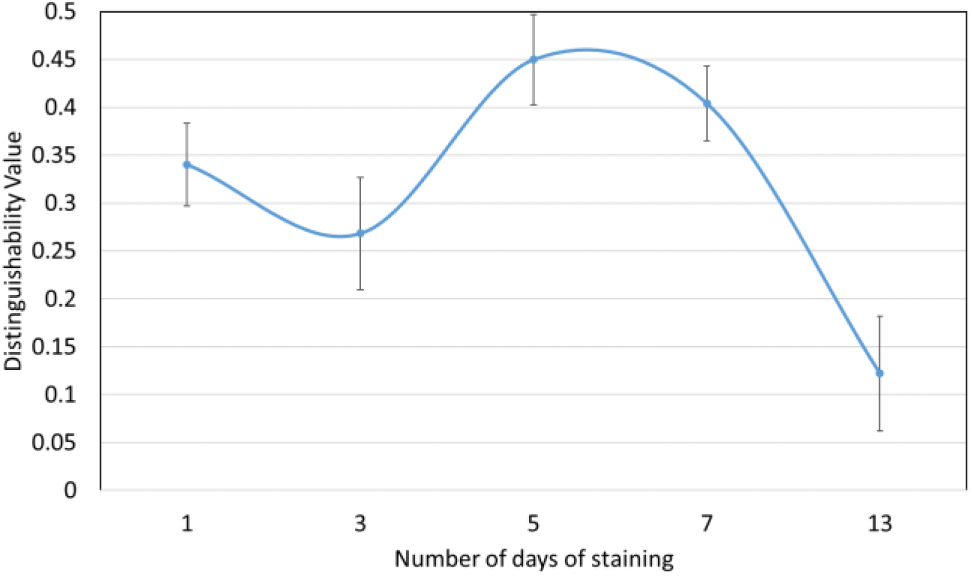
Distinguishability value for each pig vagus nerve scanned after a different number of days of staining.

The greatest distinguishability value, corresponding with the largest difference in grey value for each soft tissue type, appeared to be for the rat sciatic nerve sample stained for one day (d=0.998±0.063). Zero days of staining produced a distinguishability value of 0.063±0.070. For the pig vagus nerve, staining with iodine for five days resulted in the highest distinguishability value (d=0.449±0.047). These objective results supported what was evident after analysis by eye.

### 3.3 Optimisation of scanning parameters

The clearest tissue separation and least noise was evident with the settings used for Sample C (Figure 9). Movement artefacts were present in a few of the scans, however, calculation of SNR was performed in regions of homogeneity, avoiding the artefact, within the ROI.

**Figure 8.**
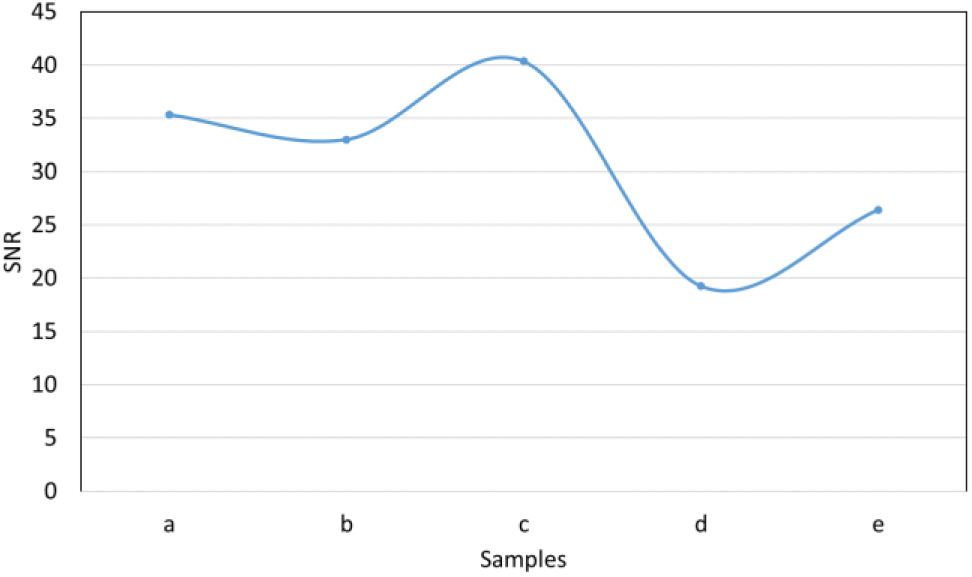
SNR value for each rat sciatic nerve sample scanned with the different parameters (Table 2).

**Figure 9.**
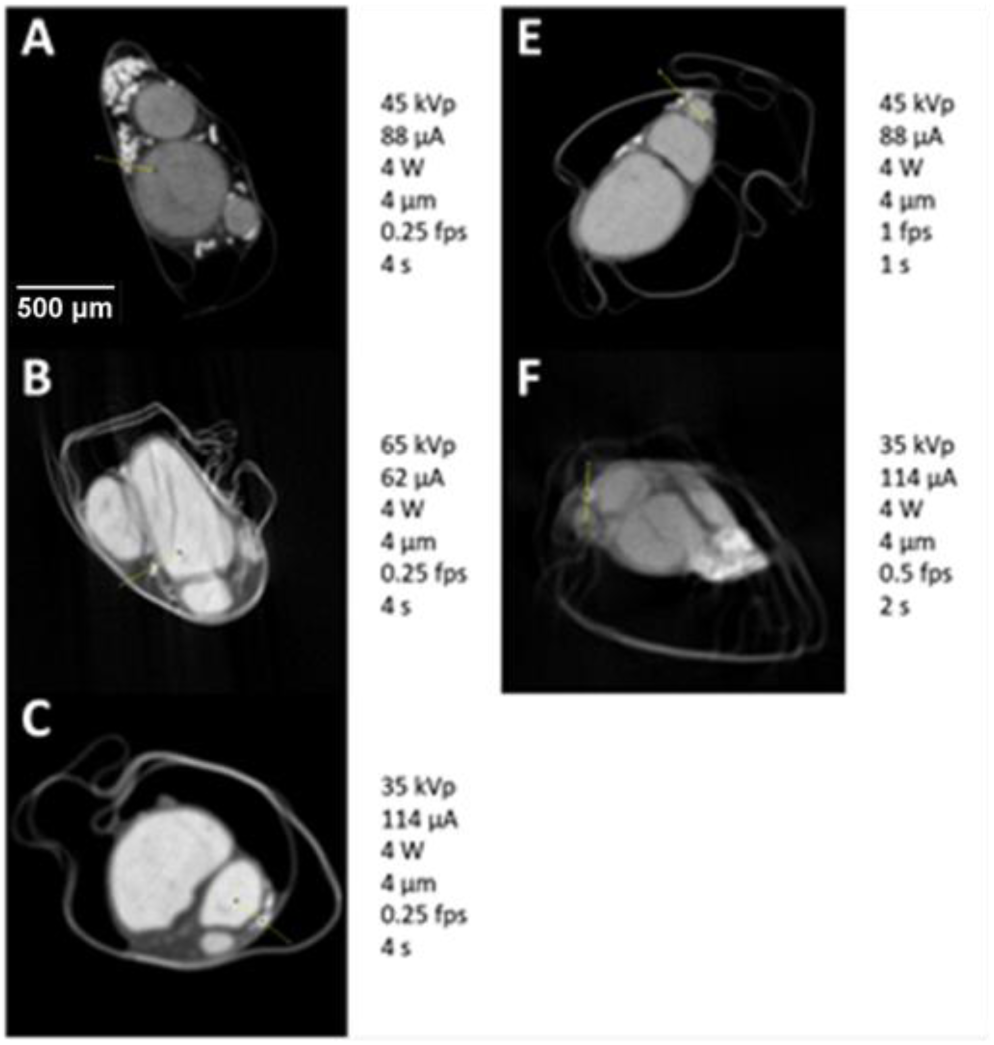
Representative XY plane slice from each scan with specific scanning parameters (listed next to the corresponding slice). Movement artefacts are present in B and F.

The image quality for each scan was quantified by calculating the SNR of fascicle-to-background for images from each scan (Figure 8). The highest SNR value (40.363) was obtained from Sample C scanned with 35 kVp energy, 114 µA current, and an exposure of 0.25 fps. This, again, validated what was evident by eye.

### 3.4 Histology for validation

Staining with H&E allowed for visualisation of all connective tissue in the nerve with the fascicles, adipocytes and connective tissue being clearly visible (Figure 10: 1.2, 2.2 and 3.2). Evident by eye, histology confirmed the positions and proportions of the corresponding fascicles in the microCT images (Figure 10, Figure 12: A and B). The average difference of the diameters of the fascicles and the full nerve between microCT and histology was 2.21%±0.0256 for rat sciatic and 2.01%±0.0237 for pig vagus nerves.

**Figure 10.**
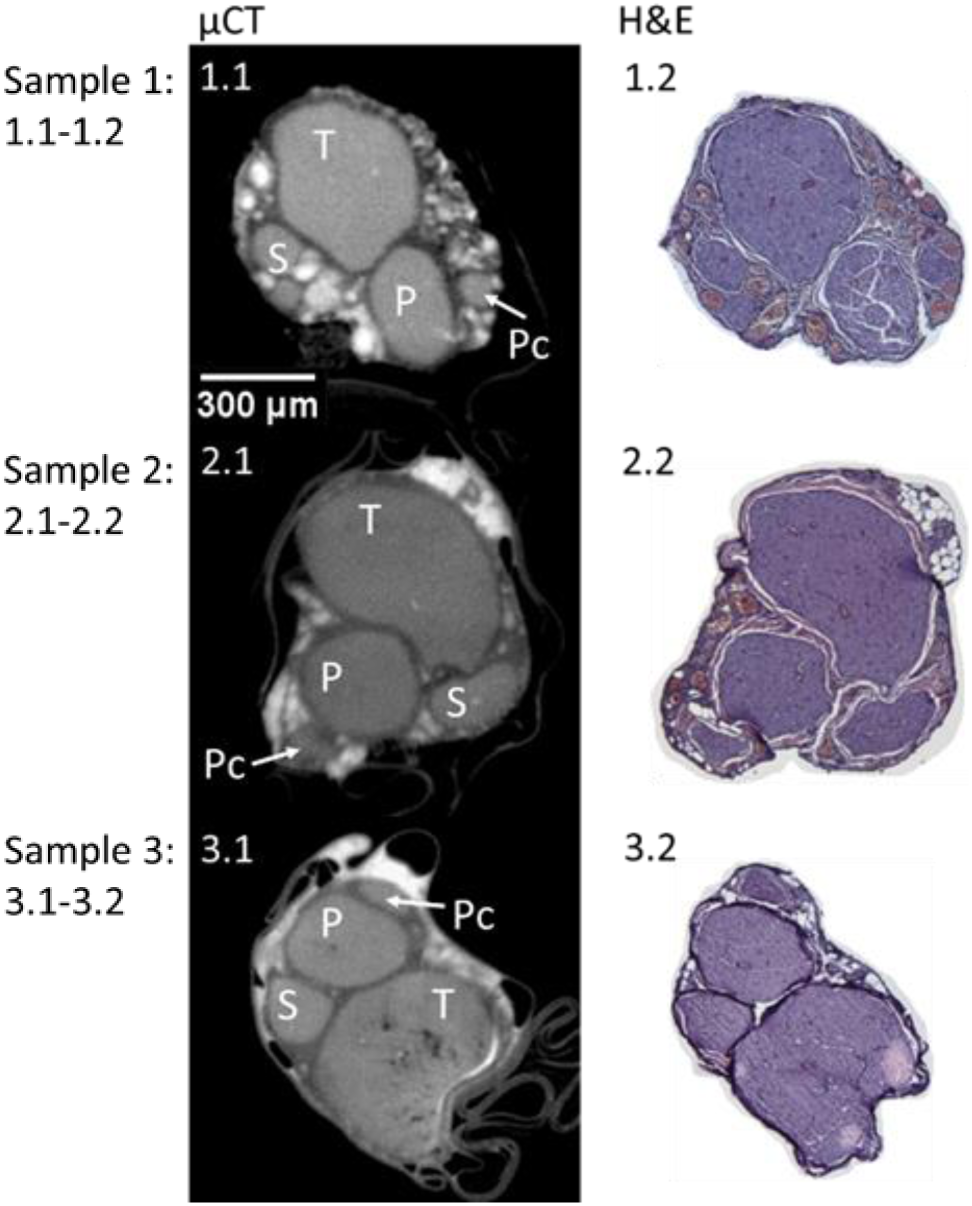
MicroCT and H&E histology slices for three example rat sciatic nerve samples. T = tibial, P = peroneal, S = sural, and Pc = post-cutaneous fascicles. n=3/12.

**Figure 11.**
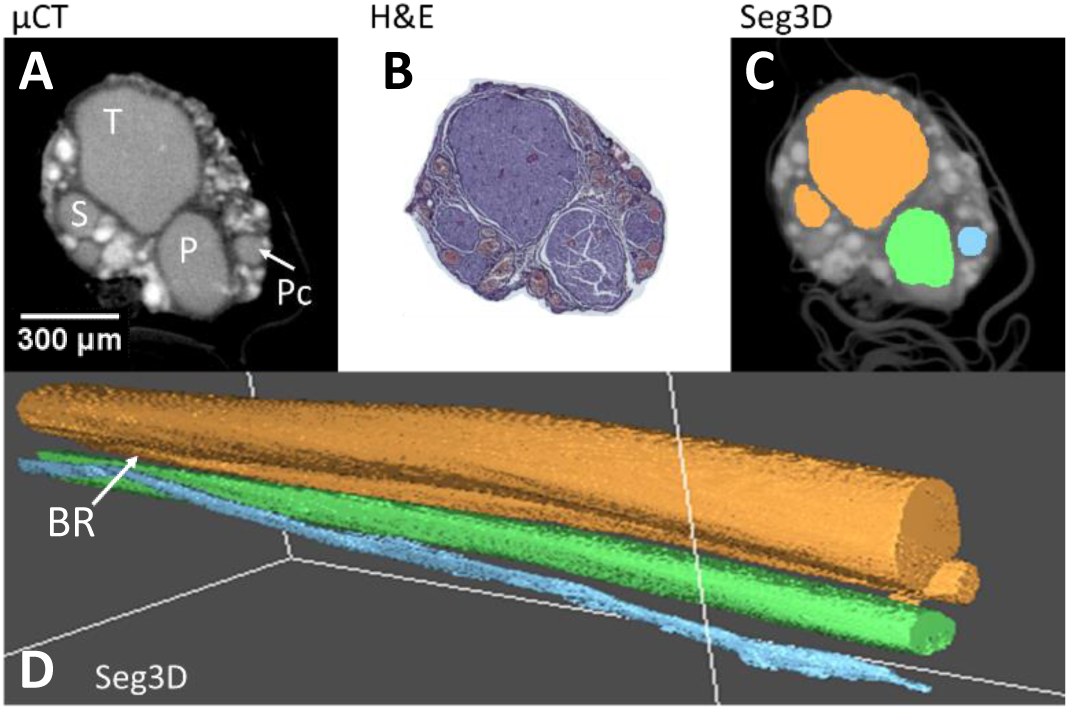
An example of a rat sciatic nerve segmentation (C, D), with its corresponding microCT using the optimal protocol (A) and histology (B). T = tibial, P = peroneal, S = sural, and Pc = post-cutaneous fascicles, and BR = branching region. n=1/12.

**Figure 12.**
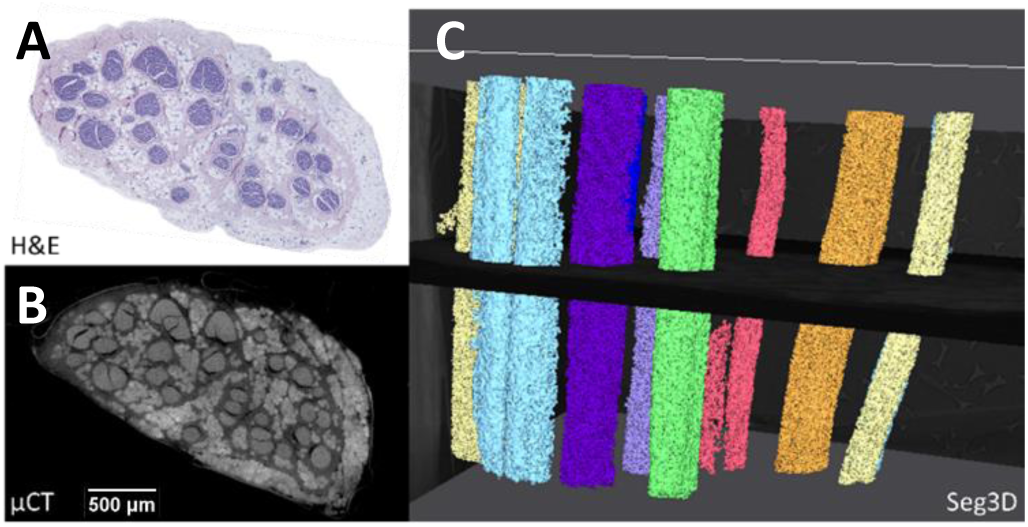
An example of a pig vagus nerve segmentation (C), with its corresponding microCT (B) and histology (A). n=1/3.

### 3.5 Computerised 3D reconstruction and fascicle tracking

Segmentation of the Lugol’s iodine-stained nerves scanned with the optimal protocol was acceptable (rat sciatic: Figure 11, pig vagus: Figure 12). Segmentation fell within the criteria for acceptability of segmenting more than three and more than ten fascicles of the rat sciatic and pig vagus nerves, respectively. In the rat sciatic nerve, for which segmentation was acceptable for four fascicles, the tibial, peroneal, sural and post-cutaneous fascicles, and a branching region between tibial and sural fascicles were present (Figure 11). In the pig vagus nerves, ten fascicles or more were able to be segmented across all scans (Figure 12). Occasionally, segmentation of one fascicle led to the segmentation of another, allowing for the discovery of branching region or a plexus between the two fascicles which was still deemed acceptable as segmentation was still indeed accurate; this often occurred if the common region (or one large fascicle), consisting and prior to branching of another two or three fascicles, was present in the scan. With minor adjustments to the Matlab script to adjust for threshold differences between scans, all scans were able to be segmented acceptably.

## 4. Discussion and Conclusions

### 4.1. Summary of results

#### Optimal protocol

Allowing for the highest differentiation between soft tissue types and thus visualisation of the ROI, the fascicles, the optimal staining time was 24 hours for rat sciatic nerves and five days for pig vagus nerves in 1% Lugol’s solution. The optimal scanning parameters that maximised the contrast between soft tissues and minimised noise and blurriness were: 35 kVp energy, 114 µA current, and an exposure of 0.25 fps. For all scans attempted, the parameters that were kept constant were: a molybdenum target, a power of 4 W, 3176 projections and a resolution with isotropic voxel size of 4 µm. With the optimal parameters, fascicles could be reconstructed and segmented acceptably over the length of the scan and the nerve.

#### How well did it work?

Histology confirmed the accuracy of fascicles’ proportions and positions from microCT to 2.21%±0.0256 for rat sciatic and 2.01%±0.0237 for pig vagus nerves. The protocol was applied to multiple nerves (rat sciatic n=12, pig vagus n=3) and was proved to be reproducible and optimised for our purposes of tracking the fascicles throughout the nerve. The distinguishability between the three soft tissue types, with specific interest in the fascicles, was successfully present in each scan and was constant throughout the length of the scans and thus, the fascicles were segmentable throughout the length of the scan and the nerve.

### 4.2. Technical issues

In early studies, there were artefacts from the movement of the nerve during the prolonged scans, due to shrinkage of the nerve from drying out over time within an open tube, used to maintain position and alignment. This was obviated by placement of the nerve in cling film within the tube to maintain as much moisture as possible. Movement artefacts were still evident. These were then corrected by 3D printing a mount that allowed for the addition of sponge around the cling film-wrapped nerve to reduce any shifting of the nerve during the scan due to gravity or rotation. Subsequent to this, the movement artefact disappeared from the images of the nerves scanned with the optimised protocol. Further evaluation criteria could possibly have been used as confirmation of the results produced, however for our purposes and after trial and error segmentation, our evaluation methods proved feasible and effective. The segmentation algorithm is semi-automatic and required manual intervention. This needs to be improved in future studies.

### 4.3. Explanations for optimal protocol

For successful segmentation of the ROI, the fascicles needed to have a distinct greyscale value (i.e. contrast) of the desired structure to the rest of the surrounding medium within the nerve including the interfascicular epineurium and adipocytes. Most segmentation methods work by identifying all the pixels, and those connected, that are within a certain data value range identified as being a part of the ROI. Staining with higher ratios or concentrations of Lugol’s results in greater degree of tissue shrinkage (Degenhardt et al., 2010). Staining with Lugol’s at a low concentration (1%) for a longer period of time, or until optimal saturation, avoids this whilst exhibiting good contrast for all tissues against background grey values (Swart et al., 2016).

For scanning, the optimal parameters produced the highest SNR and visibly had the sharpest image compared to the blurriness, noise and decreased signal seen when scanned with other parameters. It is suggested that biological samples are scanned with 30 to 100 kVp energy and in general small and low-absorbing samples, such as the rat sciatic nerve, require low voltage (du Plessis et al., 2017). To decrease the brightness of the image, thereby improving the contrast visible in the stained nerve, the voltage should be decreased, followed by the current (Sharir et al., 2011). At the same time, higher spatial resolution and statistical considerations require higher exposures to maintain the SNR (Lin et al., 2006). The highest exposure (0.25 fps) of the microCT scanner was used in the optimal protocol. Additionally, the pixel resolution used was close to the maximal pixel resolution of the scanner (3 μm per pixel). Soft tissues have a low absorption coefficient and separating the soft phases is more difficult than those of hard tissue or metal. The target selected for use with soft tissues should have a spectrum in a similar range; a material with low atomic number such as molybdenum. During reconstruction of the scans, minimal beam hardening was required to improve the image; owing to sufficient penetration of the sample (du Plessis et al., 2017).

### 4.4. Advantages over other methods

The golden standard of histology matched the microCT with high accuracy when compared at the corresponding slice, which validates the accuracy of microCT scanning and its applicability for imaging the nerves with the purpose of tracing fascicles. Tissue processing and staining for histology, for validation purposes alone, takes a minimum of three days, excluding fixation and sectioning. Therefore, in order to perform serial histology on the full length of the nerve would take a considerable amount of time longer than the two days required for microCT scanning of the entirety of the nerve. Due to limited spatial image resolution, however, the individual fibres that can be seen and analysed under the microscope are not visible in the microCT scan. Although, this is not required for our purposes of fascicle tracking. Nonetheless, our method allows for subsequent histology thereby allowing for analyses of fibres if required.

The destructiveness of the contrast agents, pre-processing methods and scanning protocols should be reduced as much as possible to ensure that the tissue integrity is not harmed (de Bournonville et al., 2019). The cost, difficult access to, fluctuations that can be attributed to other imaging techniques such as phase-contrast microCT scanning (Goyens et al., 2017) are often a limiting factor to studies with similar goals. In addition, the size of the data obtained, and thus memory required, from this more accessible Nikon scanner is significantly lower (2 to <5 GB per scan of 1 cm of nerve for rat sciatic and pig vagus nerves respectively) than from the synchrotron scanners. This obviates the requirement for supercomputers whilst still maintaining the advantage of microCT over the predominant soft-tissue imaging technique of magnetic resonance imaging (MRI) by obtaining high voxel resolution in 3D (Zhu et al., 2016). Additionally, microCT exceeds the imaging penetration depth of another prevailing imaging technique, optical coherence tomography (Islam et al., 2012).

Our optimised protocol of tissue processing, sample preparation and scanning with microCT is a simplified and reproducible method for imaging fascicular anatomy of nerves at high resolution in the axial direction whilst being time-efficient, and reducing cost, memory required and destruction of the nerve samples.

### 4.5 Recommended uses and future work

Our recommendation for the microCT imaging of the vagus and similar peripheral nerves is the staining with 1% Lugol’s solution for five days per 4 cm of nerve, wrapping in cling film and mounting in sponge within a tube, and scanning at 35 kVp energy, 114 µA current, 4 W power and an exposure of 0.25 fps, with a molybdenum target. This simplified method for imaging fascicles could not only be used to decipher the fascicular anatomy of the vagus nerve for selective stimulation but could be integrated into studies on all peripheral nerves to study peripheral nerve repair, microsurgery or improving the implementation of nerve guidance conduits for which advanced knowledge of the fascicular anatomy could prove helpful (Grinsell and Keating, 2014; Pyatin et al., 2017; Urbaniak, 1982).

The next steps will be to use the optimised protocol on the entirety of the pig vagus nerve, with its branches; scanning from the innervated organ to the cervical level and creating a fascicular map or atlas at the level of cuff placement to guide the selective VNS. Challenges that need to be overcome is extracting the entire length of the vagus nerve successfully and stitching multiple non-overlapping scans of different sections and branches of the nerve. This will eventually progress to humans to allow for improvement of VNS and its therapeutic efficiency.

## Acknowledgements

This work is supported by the Medical Research Council (MR/R01213X/1). Thank you to David Goodwin and Annette Lane, RVC, for training and assistance with histology.

## References

Amann, J.F., Constantinescu, G.M., 1990. The anatomy of the visceral and autonomic nervous systems. Semin. Vet. Med. Surg. (Small Anim.) 5, 4–11.

Aristovich, K., Donega, M., Fjordbakk, C., Tarotin, I., Chapman, C., Viscasillas, J., Stathopoulou, T.-R., Crawford, A., Chew, D., Perkins, J., Holder, D., 2019. Complete optimisation and in-vivo validation of the spatially selective multielectode array for vagus nerve neuromodulation. 190312459 Phys.

Binnie, C.D., 2000. Vagus nerve stimulation for epilepsy: a review. Seizure 9, 161–169. https://doi.org/10.1053/seiz.1999.0354

Blount, J.P., 2015. Vagus Nerve Stimulation, in: Nerves and Nerve Injuries. Elsevier, pp. 393–406. https://doi.org/10.1016/B978-0-12-802653-3.00075-0

Boerckel, J.D., Mason, D.E., McDermott, A.M., Alsberg, E., 2014. Microcomputed tomography: approaches and applications in bioengineering. Stem Cell Res. Ther. 5. https://doi.org/10.1186/scrt534

Bonaz, B., Bazin, T., Pellissier, S., 2018. The Vagus Nerve at the Interface of the Microbiota-Gut-Brain Axis. Front. Neurosci. 12. https://doi.org/10.3389/fnins.2018.00049

Breit, S., Kupferberg, A., Rogler, G., Hasler, G., 2018. Vagus Nerve as Modulator of the Brain–Gut Axis in Psychiatric and Inflammatory Disorders. Front. Psychiatry 9. https://doi.org/10.3389/fpsyt.2018.00044

Browning, K.N., Verheijden, S., Boeckxstaens, G.E., 2017. The Vagus Nerve in Appetite Regulation, Mood, and Intestinal Inflammation. Gastroenterology 152, 730–744. https://doi.org/10.1053/j.gastro.2016.10.046

Câmara, R., Griessenauer, C.J., 2015. Anatomy of the Vagus Nerve, in: Nerves and Nerve Injuries. Elsevier, pp. 385–397. https://doi.org/10.1016/B978-0-12-410390-0.00028-7

Chatterjee, S., 2014. Artefacts in histopathology. J. Oral Maxillofac. Pathol. JOMFP 18, S111–S116. https://doi.org/10.4103/0973-029X.141346

de Boer, M., Boeker, E.B., Ramrattan, M.A., Kiewiet, J.J.S., Dijkgraaf, M.G.W., Boermeester, M.A., Lie-A-Huen, L., 2013. Adverse drug events in surgical patients: an observational multicentre study. Int. J. Clin. Pharm. 35, 744–752. https://doi.org/10.1007/s11096-013-9797-5

de Bournonville, S., Vangrunderbeeck, S., Kerckhofs, G., 2019. Contrast-Enhanced MicroCT for Virtual 3D Anatomical Pathology of Biological Tissues: A Literature Review [WWW Document]. Contrast Media Mol. Imaging. https://doi.org/10.1155/2019/8617406

de Crespigny, A., Bou-Reslan, H., Nishimura, M.C., Phillips, H., Carano, R.A.D., D’Arceuil, H.E., 2008. 3D micro-CT imaging of the postmortem brain. J. Neurosci. Methods 171, 207–213. https://doi.org/10.1016/j.jneumeth.2008.03.006

De Ferrari, G.M., Schwartz, P.J., 2011. Vagus nerve stimulation: from pre-clinical to clinical application: challenges and future directions. Heart Fail. Rev. 16, 195–203. https://doi.org/10.1007/s10741-010-9216-0

Degenhardt, K., Wright, A.C., Horng, D., Padmanabhan, A., Epstein, J.A., 2010. Rapid Three-Dimensional Phenotyping of Cardiovascular Development in Mouse Embryos by Micro-CT with Iodine Staining. Circ. Cardiovasc. Imaging 3, 314–322. https://doi.org/10.1161/CIRCIMAGING.109.918482

du Plessis, A., Broeckhoven, C., Guelpa, A., le Roux, S.G., 2017. Laboratory x-ray micro-computed tomography: a user guideline for biological samples. GigaScience 6, 1–11. https://doi.org/10.1093/gigascience/gix027

Dunlop, D., 1969. Adverse Effects of Drugs. Br. Med. J. 2, 622–623.

Ekmekçi, H., Kaptan, H., 2017. Vagus Nerve Stimulation. Open Access Maced. J. Med. Sci. 5, 391–394. https://doi.org/10.3889/oamjms.2017.056

Famm, K., Litt, B., Tracey, K.J., Boyden, E.S., Slaoui, M., 2013. A jump-start for electroceuticals: Drug discovery. Nature 496, 159–161. https://doi.org/10.1038/496159a

Felten, D.L., O’Banion, M.K., Maida, M.S., 2016. NETTER’S ATLAS OF NEUROSCIENCE, 3rd ed. Elsevier, Philadelphia, PA 19103–2899.

Gignac, P.M., Kley, N.J., 2014. Iodine-enhanced micro-CT imaging: Methodological refinements for the study of the soft-tissue anatomy of post-embryonic vertebrates. J. Exp. Zoolog. B Mol. Dev. Evol. 322, 166–176. https://doi.org/10.1002/jez.b.22561

Gignac, P.M., Kley, N.J., Clarke, J.A., Colbert, M.W., Morhardt, A.C., Cerio, D., Cost, I.N., Cox, P.G., Daza, J.D., Early, C.M., Echols, M.S., Henkelman, R.M., Herdina, A.N., Holliday, C.M., Li, Z., Mahlow, K., Merchant, S., Müller, J., Orsbon, C.P., Paluh, D.J., Thies, M.L., Tsai, H.P., Witmer, L.M., 2016. Diffusible iodine-based contrast-enhanced computed tomography (diceCT): an emerging tool for rapid, high-resolution, 3-D imaging of metazoan soft tissues. J. Anat. 228, 889–909. https://doi.org/10.1111/joa.12449

Gong, H., Xu, D., Yuan, J., Li, X., Guo, C., Peng, J., Li, Y., Schwarz, L.A., Li, A., Hu, B., Xiong, B., Sun, Q., Zhang, Y., Liu, J., Zhong, Q., Xu, T., Zeng, S., Luo, Q., 2016. High-throughput dual-colour precision imaging for brain-wide connectome with cytoarchitectonic landmarks at the cellular level. Nat. Commun. 7, 12142. https://doi.org/10.1038/ncomms12142

Goyens, J., Mancini, L., Nieuwenhove, V.V., Sijbers, J., Aerts, P., 2017. Comparison of conventional and synchrotron X-ray microCT scanning of thin membranes in the inner ear 4.

Grinsell, D., Keating, C.P., 2014. Peripheral Nerve Reconstruction after Injury: A Review of Clinical and Experimental Therapies [WWW Document]. BioMed Res. Int. https://doi.org/10.1155/2014/698256

Guiraud, D., Andreu, D., Bonnet, S., Carrault, G., Couderc, P., Hagège, A., Henry, C., Hernandez, A., Karam, N., Le Rolle, V., Mabo, P., Maciejasz, P., Malbert, C.-H., Marijon, E., Maubert, S., Picq, C., Rossel, O., Bonnet, J.-L., 2016. Vagus nerve stimulation: state of the art of stimulation and recording strategies to address autonomic function neuromodulation. J. Neural Eng. 13, 041002. https://doi.org/10.1088/1741-2560/13/4/041002

Heimel, P., Swiadek, N.V., Slezak, P., Kerbl, M., Schneider, C., Nürnberger, S., Redl, H., Teuschl, A.H., Hercher, D., 2019. Iodine-Enhanced Micro-CT Imaging of Soft Tissue on the Example of Peripheral Nerve Regeneration. Contrast Media Mol. Imaging 2019. https://doi.org/10.1155/2019/7483745

Iannaccone, P.M., Jacob, H.J., 2009. Rats! Dis. Model. Mech. 2, 206–210. https://doi.org/10.1242/dmm.002733

Islam, M.S., Oliveira, M.C., Wang, Y., Henry, F.P., Randolph, M.A., Park, B.H., de Boer, J.F., 2012. Extracting structural features of rat sciatic nerve using polarization-sensitive spectral domain optical coherence tomography. J. Biomed. Opt. 17, 056012. https://doi.org/10.1117/1.JBO.17.5.056012

Jeffery, N.S., Stephenson, R.S., Gallagher, J.A., Jarvis, J.C., Cox, P.G., 2011. Micro-computed tomography with iodine staining resolves the arrangement of muscle fibres. J. Biomech. 44, 189–192. https://doi.org/10.1016/j.jbiomech.2010.08.027

Johnson, J.T., Hansen, M.S., Wu, I., Healy, L.J., Johnson, C.R., Jones, G.M., Capecchi, M.R., Keller, C., 2006. Virtual histology of transgenic mouse embryos for high-throughput phenotyping. PLoS Genet. 2, e61. https://doi.org/10.1371/journal.pgen.0020061

Kalson, N.S., Malone, P.S.C., Bradley, R.S., Withers, P.J., Lees, V.C., 2012. Fibre bundles in the human extensor carpi ulnaris tendon are arranged in a spiral. J. Hand Surg. Eur. Vol. 37, 550–554. https://doi.org/10.1177/1753193411433228

Kaplan, H.M., Mishra, P., Kohn, J., 2015. The overwhelming use of rat models in nerve regeneration research may compromise designs of nerve guidance conduits for humans. J. Mater. Sci. Mater. Med. 26. https://doi.org/10.1007/s10856-015-5558-4

Klein, H.U., Ferrari, G.M.D., 2010. Vagus nerve stimulation: A new approach to reduce heart failure. Cardiol. J. 17, 638–644.

Kollewe, J., 2017. Electroceuticals: the “bonkers” gamble that could pay off for GlaxoSmithKline. The Guardian.

Koopman, F.A., Chavan, S.S., Miljko, S., Grazio, S., Sokolovic, S., Schuurman, P.R., Mehta, A.D., Levine, Y.A., Faltys, M., Zitnik, R., Tracey, K.J., Tak, P.P., 2016. Vagus nerve stimulation inhibits cytokine production and attenuates disease severity in rheumatoid arthritis. Proc. Natl. Acad. Sci. 113, 8284–8289. https://doi.org/10.1073/pnas.1605635113

Lin, M.D., Samei, E., Badea, C.T., Yoshizumi, T.T., Johnson, G.A., 2006. Optimized radiographic spectra for small animal digital subtraction angiography. Med. Phys. 33, 4249–4257. https://doi.org/10.1118/1.2356646

McCorry, L.K., 2007. Physiology of the Autonomic Nervous System. Am. J. Pharm. Educ. 71.

McInnes, E., 2005. Artefacts in histopathology. Comp. Clin. Pathol. 13, 100–108. https://doi.org/10.1007/s00580-004-0532-4

Metscher, B.D., 2009. MicroCT for comparative morphology: simple staining methods allow high-contrast 3D imaging of diverse non-mineralized animal tissues. BMC Physiol. 9, 11. https://doi.org/10.1186/1472-6793-9-11

Mishra, S., 2017. Electroceuticals in medicine – The brave new future. Indian Heart J. 69, 685–686. https://doi.org/10.1016/j.ihj.2017.10.001

Mizutani, R., Takeuchi, A., Hara, T., Uesugi, K., Suzuki, Y., 2007. Computed tomography imaging of the neuronal structure of Drosophila brain. J. Synchrotron Radiat. 14, 282–287. https://doi.org/10.1107/S0909049507009004

O’Sullivan, J.D.B., Behnsen, J., Starborg, T., MacDonald, A.S., Phythian-Adams, A.T., Else, K.J., Cruickshank, S.M., Withers, P.J., 2018. X-ray micro-computed tomography (μCT): an emerging opportunity in parasite imaging. Parasitology 145, 848–854. https://doi.org/10.1017/S0031182017002074

Pauwels, E., Loo, D.V., Cornillie, P., Brabant, L., Hoorebeke, L.V., 2013. An exploratory study of contrast agents for soft tissue visualization by means of high resolution X-ray computed tomography imaging. J. Microsc. 250, 21–31. https://doi.org/10.1111/jmi.12013

Pečlin, P., Rozman, J., 2014. Alternative Paradigm of Selective Vagus Nerve Stimulation Tested on an Isolated Porcine Vagus Nerve. Sci. World J. 2014, 1–10. https://doi.org/10.1155/2014/310283

Pyatin, V.F., Kolsanov, A.V., Shirolapov, I.V., 2017. Recent medical techniques for peripheral nerve repair: Clinico-physiological advantages of artificial nerve guidance conduits. Adv. Gerontol. 7, 148–154. https://doi.org/10.1134/S2079057017020126

Rea, P., 2014. Chapter 10 - Vagus Nerve, in: Rea, P. (Ed.), Clinical Anatomy of the Cranial Nerves. Academic Press, San Diego, pp. 105–116. https://doi.org/10.1016/B978-0-12-800898-0.00010-5

Ripplinger, C.M., 2017. From drugs to devices and back again: chemical vagal nerve stimulation for the treatment of heart failure. Cardiovasc. Res. 113, 1270–1272. https://doi.org/10.1093/cvr/cvx142

Schindelin, J., Arganda-Carreras, I., Frise, E., Kaynig, V., Longair, M., Pietzsch, T., Preibisch, S., Rueden, C., Saalfeld, S., Schmid, B., Tinevez, J.-Y., White, D.J., Hartenstein, V., Eliceiri, K., Tomancak, P., Cardona, A., 2012. Fiji: an open-source platform for biological-image analysis. Nat. Methods 9, 676–682. https://doi.org/10.1038/nmeth.2019

Scott, A.E., Vasilescu, D.M., Seal, K.A.D., Keyes, S.D., Mavrogordato, M.N., Hogg, J.C., Sinclair, I., Warner, J.A., Hackett, T.-L., Lackie, P.M., 2015. Three Dimensional Imaging of Paraffin Embedded Human Lung Tissue Samples by Micro-Computed Tomography. PLOS ONE 10, e0126230. https://doi.org/10.1371/journal.pone.0126230

Senter-Zapata, M., Patel, K., Bautista, P.A., Griffin, M., Michaelson, J., Yagi, Y., 2016. The Role of Micro-CT in 3D Histology Imaging. Pathobiology 83, 140–147. https://doi.org/10.1159/000442387

Sharir, A., Ramniceanu, G., Brumfeld, V., 2011. High Resolution 3D Imaging of Ex-Vivo Biological Samples by Micro CT. J. Vis. Exp. JoVE. https://doi.org/10.3791/2688

Shearer, T., Bradley, R.S., Hidalgo-Bastida, L.A., Sherratt, M.J., Cartmell, S.H., 2016. Three-dimensional visualisation of soft biological structures by X-ray computed micro-tomography. J. Cell Sci. 129, 2483–2492. https://doi.org/10.1242/jcs.179077

Shearer, T., Rawson, S., Castro, S.J., Balint, R., Bradley, R.S., Lowe, T., Vila-Comamala, J., Lee, P.D., Cartmell, S.H., 2014. X-ray computed tomography of the anterior cruciate ligament and patellar tendon. Muscles Ligaments Tendons J. 4, 238–244.

Sheehan, D.C., Hrapchak, B.B., 1987. Theory and practice of histotechnology. Battelle Press, Columbus, Ohio.

Sibilla, L., Agarwal, N., 2018. Cranial Nerve X: Vagus, in: Agarwal, N., Port, J.D. (Eds.), Neuroimaging: Anatomy Meets Function. Springer International Publishing, Cham, pp. 211–214. https://doi.org/10.1007/978-3-319-57427-1_21

Smucny, J., Visani, A., Tregellas, J.R., 2015. Could Vagus Nerve Stimulation Target Hippocampal Hyperactivity to Improve Cognition in Schizophrenia? Front. Psychiatry 6. https://doi.org/10.3389/fpsyt.2015.00043

Stauber, M., Müller, R., 2008. Micro-Computed Tomography: A Method for the Non-Destructive Evaluation of the Three-Dimensional Structure of Biological Specimens, in: Westendorf, J.J. (Ed.), Osteoporosis: Methods and Protocols, Methods In Molecular BiologyTM. Humana Press, Totowa, NJ, pp. 273–292. https://doi.org/10.1007/978-1-59745-104-8_19

Sunderland, S., 1978. Nerves and nerve injuries / Sir Sydney Sunderland; foreword by the late Sir Francis Walshe, 2nd ed. ed. Edinburgh: Churchill Livingstone, Edinburgh.

Swart, P., Wicklein, M., Sykes, D., Ahmed, F., Krapp, H.G., 2016. A quantitative comparison of micro-CT preparations in Dipteran flies. Sci. Rep. 6. https://doi.org/10.1038/srep39380

Thompson, N., Mastitskaya, S., Holder, D., 2019. Avoiding off-target effects in electrical stimulation of the cervical vagus nerve: Neuroanatomical tracing techniques to study fascicular anatomy of the vagus nerve. J. Neurosci. Methods 325, 108325. https://doi.org/10.1016/j.jneumeth.2019.108325

Urbaniak, J.R., 1982. Fascicular nerve suture. Clin. Orthop. 57–64.

Verlinden, T.J.M., Rijkers, K., Hoogland, G., Herrler, A., 2016. Morphology of the human cervical vagus nerve: implications for vagus nerve stimulation treatment. Acta Neurol. Scand. 133, 173–182. https://doi.org/10.1111/ane.12462

Wirkner, C.S., Prendini, L., 2007. Comparative morphology of the hemolymph vascular system in scorpions--a survey using corrosion casting, MicroCT, and 3D-reconstruction. J. Morphol. 268, 401–413. https://doi.org/10.1002/jmor.10512

Wirkner, C.S., Richter, S., 2004. Improvement of microanatomical research by combining corrosion casts with MicroCT and 3D reconstruction, exemplified in the circulatory organs of the woodlouse. Microsc. Res. Tech. 64, 250–254. https://doi.org/10.1002/jemt.20076

Wolthuis, A.M., Stakenborg, N., D’Hoore, A., Boeckxstaens, G.E., 2016. The pig as preclinical model for laparoscopic vagus nerve stimulation. Int. J. Colorectal Dis. 31, 211–215. https://doi.org/10.1007/s00384-015-2435-z

XT H 225 | Computed Tomography | X-ray and CT inspection | Nikon Metrology [WWW Document], n.d. URL https://www.nikonmetrology.com/en-gb/product/xt-h-225 (accessed 6.27.19).

Yan, L., Guo, Y., Qi, J., Zhu, Q., Gu, L., Zheng, C., Lin, T., Lu, Yutong, Zeng, Z., Yu, S., Zhu, S., Zhou, X., Zhang, X., Du, Y., Yao, Z., Lu, Yao, Liu, X., 2017. Iodine and freeze-drying enhanced high-resolution MicroCT imaging for reconstructing 3D intraneural topography of human peripheral nerve fascicles. J. Neurosci. Methods 287, 58–67. https://doi.org/10.1016/j.jneumeth.2017.06.009

Zhu, S., Zhu, Q., Liu, X., Yang, W., Jian, Y., Zhou, X., He, B., Gu, L., Yan, L., Lin, T., Xiang, J., Qi, J., 2016. Three-dimensional Reconstruction of the Microstructure of Human Acellular Nerve Allograft. Sci. Rep. 6, 30694. https://doi.org/10.1038/srep30694

Zimmerman, H.J., 1999. Hepatotoxicity: The Adverse Effects of Drugs and Other Chemicals on the Liver. Lippincott Williams & Wilkins.

